# Human sensorimotor cortex reactivates recent visuomotor experience during awake rest

**DOI:** 10.1101/2024.05.26.595974

**Authors:** Kenji Ogawa, Yuxiang Yang, Huixiang Yang, Fumihito Imai, Hiroshi Imamizu

## Abstract

Previous studies have suggested that awake rest after training is helpful in improving motor performance and memory consolidation in visuomotor learning. Re-emergence of task-related activation patterns during awake rest has been reported, which play a role in memory consolidation or perceptual learning. This study aimed to test whether such reactivation occurs after visuomotor learning in the primary sensorimotor cortex. During fMRI scanning, 42 normal participants learned visuomotor tracking, while a rotational perturbation was introduced between a cursor position and a joystick angle. This visuomotor learning block was interleaved with the control block, during which the participants passively viewed a replay of previously performed cursor movements of their own. Half of the participants used their right hand, and the other half used their left hand to control the joystick. The resting-state scans were measured before and after the visuomotor learning sessions. A multivariate pattern classifier was trained to classify task and control blocks and then tested with resting scans before and after learning. Results revealed a significant increase in the number of volumes classified as the task in the post-learning rest compared with the pre-learning, indicating a re-emergence of task-related activities. Representational similarity analysis also showed a more similar pattern of activity with the task during the post-learning rest period. Furthermore, this effect is specific to the primary sensorimotor cortex contralateral to the hand used and significantly correlated with motor improvement after rest. Our finding revealed the reactivation of task-related patterns in the primary sensorimotor cortex for offline visuomotor learning.

**Significance Statement:** Previous research suggests that awake rest after learning promotes memory consolidation, which is subserved by the re-emergence of task-specific activity patterns. We aimed to determine whether such reactivation occurs in the primary sensorimotor cortex following visuomotor learning for offline memory consolidation. Our results showed a significant increase in task-classified brain volumes during the post-learning rest period compared to the pre-learning period, indicating a re-emergence of task-related activity. Furthermore, this effect was specific to the primary sensorimotor cortex contralateral to the hand used for the task and significantly correlated with the motor performance following the rest period. These findings provide evidence for the reactivation of task-related patterns during offline visuomotor learning, which may underlie memory consolidation processes.

## Introduction

Humans can flexibly acquire various motor skills through learning, and such motor memories are stored as internal models in the central nervous system (Imamizu et al., 2000; Wolpert and Ghahramani, 2000). In visuomotor learning, previous studies have suggested that sleep as well as awake rest after learning is helpful in improving behavioral performance and memory consolidation (Brashers-Krug et al., 1996; Shadmehr and Brashers-Krug, 1997; Robertson et al., 2004). For example, motor performance could be improved after a period of rest rather than immediately training a new skill, and motor memory is more consolidated and less vulnerable to interference from new motor skills (Robertson et al., 2004, 2005; Cohen et al., 2005; Press et al., 2005). It has also been reported that introducing periods of waking rest between learning sessions has a positive effect on the retention of motor memory, known as the spacing effect (Cepeda et al., 2006; Kornmeier and Sosic-Vasic, 2012; Gerbier et al., 2015). These studies suggest that the waking rest period after learning plays an essential role in the offline learning of motor memory.

The brain generates activity spontaneously, even when no specific task is required (Raichle et al., 2001; Fox and Raichle, 2007). It has been shown that spontaneous activation during rest is modulated by perceptual or motor learning (Tambini and Davachi, 2019). Animal studies suggested that resting-state brain activity represents a prior distribution in visual perception, constructing an internal model of the environment (Berkes et al., 2011). Human fMRI studies have shown that the resting-state activity in the early visual cortex changes according to visual perceptual learning (Lewis et al., 2009; Guidotti et al., 2015) or enhanced resting-state functional connectivity between the hippocampus and a portion of the lateral occipital complex related to memory consolidation (Tambini et al., 2010). In motor learning, human studies have reported a widespread enhanced activation of sensorimotor networks during awake rest following training with fMRI (Albert et al., 2009; Vahdat et al., 2011; Sami et al., 2014; Lin et al., 2018) or EEG (Wu et al., 2014; Gentili et al., 2015), with a link to improvement through offline learning (Gregory et al., 2014; Manuel et al., 2018).

Neural replay during rest after training is thought to be a mechanism for consolidating memory, reflecting cortical plasticity (Kurth-Nelson et al., 2023). Replay or reactivation in the hippocampus is considered an active system of memory consolidation (Klinzing et al., 2019). The first empirical evidence for this phenomenon was found by recordings of neuronal assemblies of rodents during sleep, which showed that the hippocampus exhibits autonomous reactivation of neuronal assemblies that were engaged during the previous experiential episodes (Pavlides and Winson, 1989). Subsequent studies have shown that a similar ‘replay’ or reactivation phenomenon occurs in the visual cortex of rodents during sleep (Ji and Wilson, 2007) as well as the awake resting period (Han et al., 2008). The autonomic reactivation lasting several minutes after repetitive visual stimulus is thought to facilitate short-term memory and contribute to long-term perceptual learning (Han et al., 2008). However, it remains to be elucidated whether there are reactivations of neural populations related to learned sensorimotor skills, together with the relationship to offline improvement in behavioral performance.

This study aimed to test whether such re-emergence of activation occurs after visuomotor learning in the human sensorimotor cortex. In our experiments, the participants performed continuous manual visuomotor tracking within an MRI scanner, and we measured brain activations with fMRI during both the learning period and the rest period before and after the learning. Using a multi-voxel pattern analysis (MVPA), we first tried to test whether brain activity patterns are similar to those during the previous motor task (reactivation) in the resting period. Furthermore, we tested whether such reactivation has a facilitatory effect on behavioral performance after learning.

## Materials and Methods

### Participants

Participants included 42 volunteers (29 males and 13 females) with a mean age of 22.7 years (range, 20–34 years). Sample size was determined before data collection based on our previous study, which used MVPA with the two groups of participants to compare the neural representation of the primary sensorimotor cortex (Ogawa et al., 2019). Half of the participants (n = 21, 6 female) used their right hand, and the other half (n = 21, 7 female) used their left hand to control the joystick in the MRI scanner. One participant (male) of the left-hand group was excluded from the analysis because he did not move the cursor frequently during the task period. All participants were right-handed, as assessed by a modified version of the Edinburgh Handedness Inventory (Oldfield, 1971) modified for Japanese participants (Hatta and Nakatsuka, 1975). Written informed consent was obtained from all participants in accordance with the Declaration of Helsinki. The experimental protocol received approval from the local ethics committee.

### Task procedures

The participants underwent the fMRI scanning, which consisted of 4 task sessions and 2 resting-state (RS) scanning sessions before (pre-RS) and after (post-RS) the first 3 task sessions (Figure 1A). In the task sessions, the participants performed continuous visuomotor tracking movement (Ogawa and Imamizu, 2013), while a rotational perturbation of 30° was introduced between a cursor position and a joystick angle. In the visuomotor learning block (task block, 12 s), the participants were instructed to chase a randomly moving target in the frame by controlling the cursor with the joystick. This visuomotor learning block was interleaved with the replay block (12 s), during which the participants passively viewed a replay of previously performed cursor movements of their own. The task session consisted of 10 task blocks and 10 replay blocks. In the RS sessions, participants were asked to stare at the cross in the center, not to move their bodies, not to sleep, and to remain rested. Each RS session lasted 6 minutes (Figure 1B). Finally, the participants underwent a T1 anatomical scanning.

**Figure 1:**
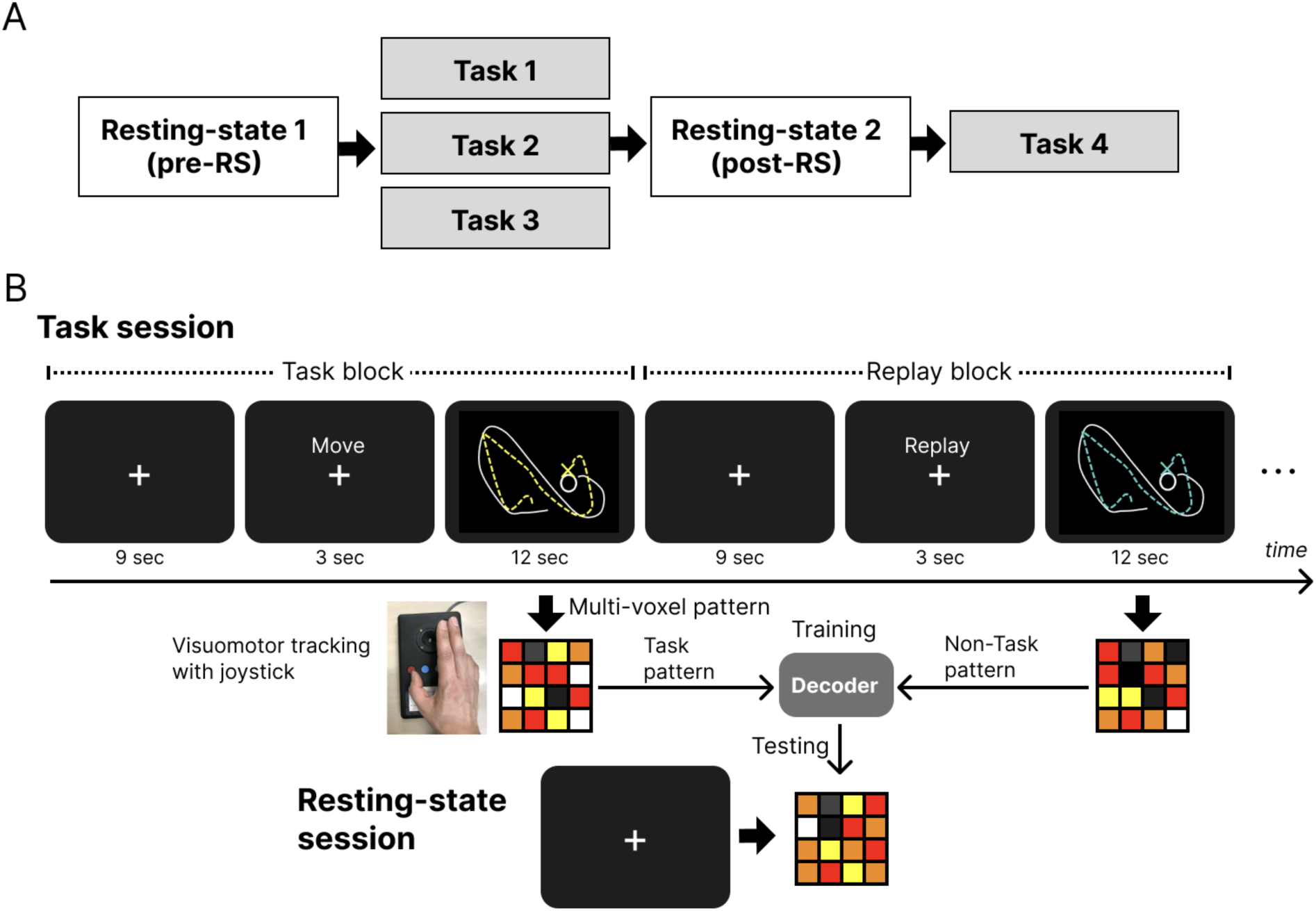
Schematic description of experimental time-course and multi-voxel pattern analysis (MVPA) A) Experimental schedule of the task and resting-scan sessions. B) The upper part shows the example time-course for one trial consisting of 1 task block and 1 replay block within the task session. The lower part shows the activity patterns during the task and replay blocks within the task sessions, which are used as task and non-task patterns for training of the decoder. The task and non-task patterns in 3 task sessions are used to train the decoder and then tested the pre-RS and post-RS activities.

### MRI acquisition

All scans were performed on a Siemens (Erlangen, Germany) 3-Tesla Prisma scanner with a 64-channel head coil at Hokkaido University. T2*-weighted echo-planar imaging (EPI) was used to acquire a total of 174 scans per task session and 122 scans per rest session, with a gradient EPI sequence. The first three scans within each session were discarded to allow for T1 equilibration. The scanning parameters were repetition time (TR), 3000 ms; echo time (TE), 30 ms; flip angle (FA), 90°; field of view (FOV), 192 × 192 mm; matrix, 94 × 94; 36 axial slices; and slice thickness, 3.0 mm with a 0.75 mm gap. T1-weighted anatomical imaging with an MP-RAGE sequence was performed using the following parameters: TR, 2300 ms; TE, 2.41 ms; FA, 8°; FOV; 256 × 256 mm; matrix, 256 × 256; 224 axial slices; and slice thickness, 0.8 mm without a gap.

### fMRI mass-univariate analysis

Image preprocessing was performed using the SPM12 software (Welcome Department of Cognitive Neurology, http://www.fil.ion.ucl.ac.uk/spm). All functional images were initially realigned to adjust for motion-related artifacts. Volume-based realignment was performed by co-registering images using rigid body transformation to minimize the squared differences between volumes. The realigned images were then spatially normalized with the Montreal Neurological Institute template based on the affine and nonlinear registration of coregistered T1-weighted anatomical images (normalization procedure of SPM). They were resampled into 3-mm-cube voxels with the sinc interpolation. Images were spatially smoothed using a Gaussian kernel of 6 × 6 × 6 mm full width at half-maximum. However, images used for MVPA were not smoothed to avoid blurring the fine-grained information contained in the multivoxel activity (Mur et al., 2009; Kamitani and Sawahata, 2010). We analyzed significantly activated areas during the task block compared with the replay (observation only) block with the mass univoxel analysis. Activation was the threshold at p < .05 corrected for multiple comparisons for a family-wise error (FWE), with an extent threshold of 15 voxels.

### MVPA

We used MVPA to classify the task and replay activities using a spatiotemporal decoder (Guidotti et al., 2015). The classifier was based on a linear support vector machine run by LIBSVM (http://www.csie.ntu.edu.tw/~cjlin/libsvm) with a fixed regularization parameter C = 1. The region of interest (ROI) was defined anatomically with the precentral and postcentral cortices of the automated anatomical labeling (AAL) toolbox (Tzourio-Mazoyer et al., 2002) as the primary sensorimotor cortex. First, we tried to classify the brain activities of the task block and those of the replay block with leave one-session out cross-validation among the first 3 task sessions. The activation pattern during the task block and those during the replay block with 4 consecutive volumes in the 3 task sessions were used as the spatio-temporal patterns to train the decoder (Figure 1B middle). This analysis produced the mean classification accuracy among the 3 task sessions using leave one-session-out cross-validation. Next, we investigated whether the reappearance of task-related activation patterns occurred in the primary sensorimotor cortex more frequently after task training (post-RS) than before the task period (pre-RS). As in the above analysis of the task sessions, the decoder was first trained to classify the brain activities between the task block and the replay blocks using the 3 task sessions as the training data. This decoder was then tested with the activities of the pre-RS and post-RS to see whether a task-like brain activity pattern occurred during the rest session (Figure 1B bottom). This analysis used a sliding time window, where a window the length of the task or replay block (4 volumes) was slid through the resting scan volumes, advancing one volume per analysis.

Pattern similarity-based classification was also conducted using representational similarity analysis (RSA) (Kriegeskorte et al., 2008). We calculated the Euclidean distance between brain activation patterns during pre- and post-rest and task/non-task patterns. If the task pattern was close to the sample in the test dataset, this data was classified as a task pattern; otherwise, it was classified as a non-task (observation of replay) pattern.

## Results

### Behavioral results

We conducted a mixed-effects analysis of variance (ANOVA) on the average tracking error between the right-hand and left-hand groups during the four task sessions. The results showed a significant main effect of the groups (*F*(1, 39) = 7.65, *p* = .01, *η_p_^2^* = .16) and of the sessions (*F*(3, 117) = 17.19, *p* < .001, *η_p_^2^* = .31) with no significant interaction (*F*(3, 117) = 1.16, *p* = .33, *η_p_^2^* = .03). Post-hoc comparisons showed a significantly larger tracking error in task session 1 compared with session 2 (*t*(40) = 4.46, *p_Bonf_* < .001), session 3 (*t*(40) = 4.75, *p_Bonf_* < .001) and session 4 (*t*(40) = 7.01, *p_Bonf_* < .001) with a marginally significant difference between session 2 and 4 (*t*(40) = 2.55, *p_Bonf_* < .1) (Figure 2).

**Figure 2:**
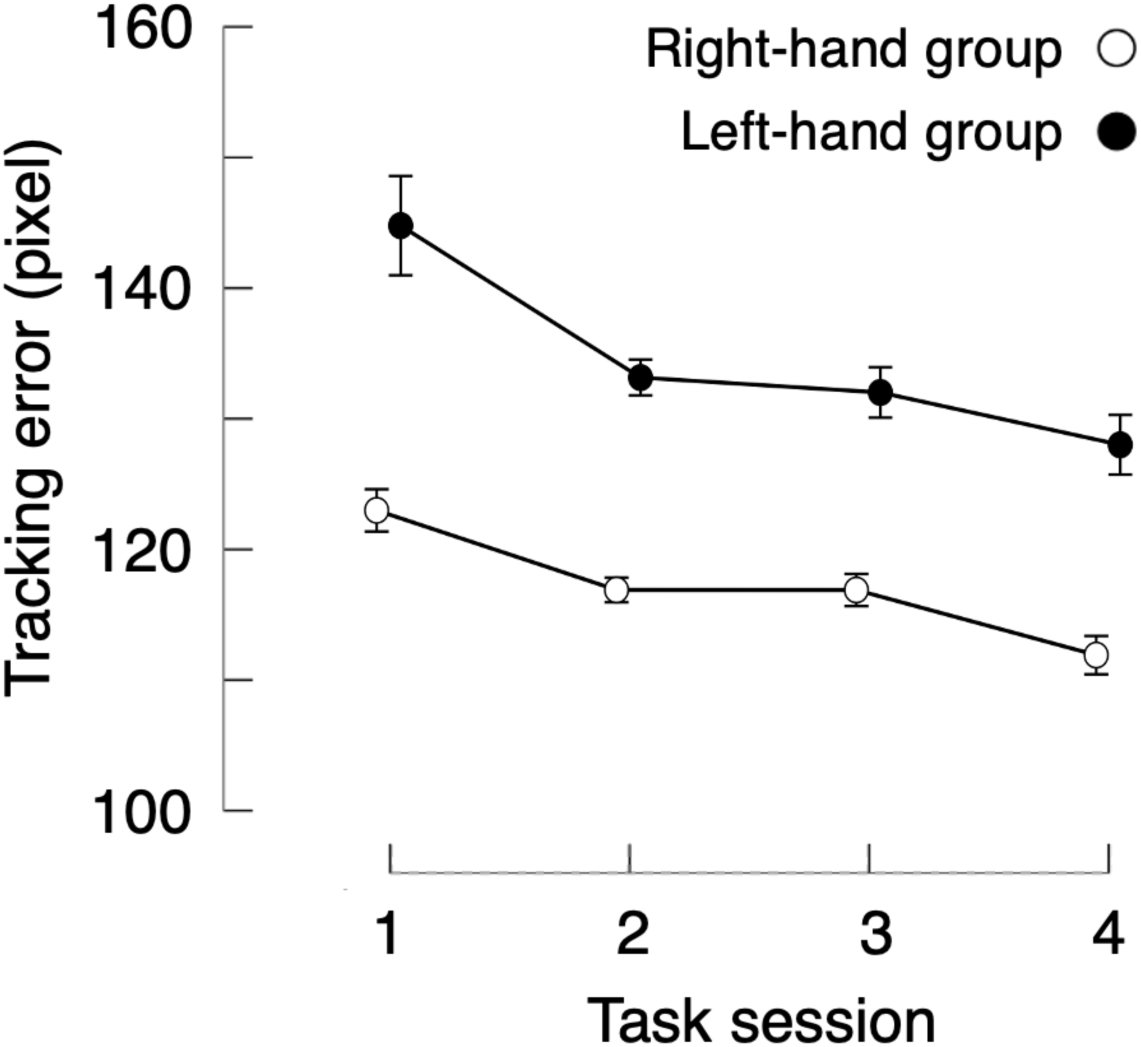
Behavioral results of tracking error The average tracking error between the target and the cursor (in pixels) during the task sessions for the right-hand group (white dots) and the left-hand group (black dots). The error bars show SEMs.

### fMRI mass-univariate analysis

We analyzed the activated regions of the brain using the conventional mass-univariate analysis of single voxels for each group. We compared the activities between the task blocks and replay blocks within four task sessions to reveal the brain regions activated during the task block. We then found the activations mainly in the primary sensorimotor cortex and the cerebellum, which are either contralateral or ipsilateral to the hand used, together with the small clusters in the thalamus, the basal ganglia, and the central operculum (Figure 3; Table 1).

**Figure 3:**
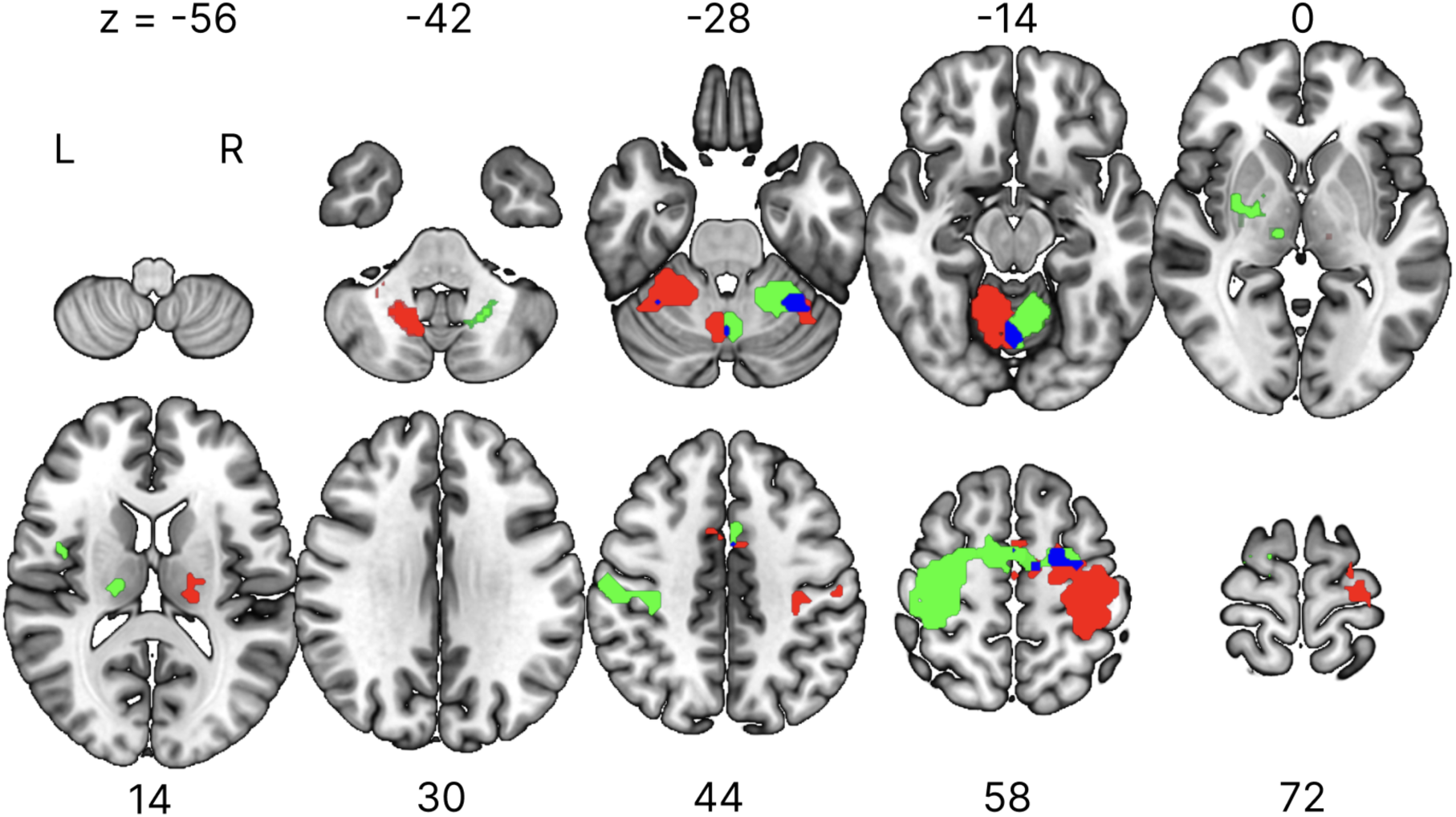
Activated areas during the task block with the mass-univariate analysis. Areas activated during the task block compared with the replay block in task sessions for the right-hand group (green) and the left-hand group (red) with their overlap (blue). Activation was reported with a threshold of p < .05 corrected for multiple comparisons for family-wise error (FWE) with an extent threshold of 15 voxels. MNI coordinates of activated foci are reported in Table 1. The regions are displayed in the horizontal plane with Z denoting locations in the MNI coordinates. L, left hemisphere; R, right hemisphere.

**Table 1:** Anatomical regions, peak voxel coordinates, and t-values of observed activation during task compared with replay blocks for each group.

| Anatomic region | voxels | MNI coordinates |  |  | *t*-value |
| --- | --- | --- | --- | --- | --- |
|  |  | x | y | z |  |
| *Right-hand group* |  |  |  |  |  |
| R cerebellum | 673 | 21 | -49 | -22 | 20.45 |
| L precentral/postcentral gyrus | 713 | -30 | -25 | 56 | 16.09 |
| L thalamus | 55 | -12 | -19 | 5 | 11.47 |
| L cerebellum | 63 | -30 | -58 | -25 | 5.62 |
| *Left-hand group* |  |  |  |  |  |
| L cerebellum | 400 | -18 | -52 | -22 | 16.35 |
| R precentral/postcentral gyrus | 845 | 36 | -13 | 59 | 15.39 |
| R thalamus | 55 | 12 | -19 | 5 | 15.08 |
| L cerebellum | 62 | -21 | -58 | -46 | 12.43 |
| R basal ganglia | 38 | 21 | -7 | -1 | 5.94 |
| R central operculum | 22 | 42 | -1 | 14 | 9.66 |
| R cerebellum | 15 | 30 | -55 | -22 | 5.30 |

Activation was reported with a threshold of $p < 0.05$ corrected for family-wise error (FWE) with an extent threshold of 15 voxels. MNI, Montreal Neurological Institute; L, left hemisphere; R, right hemisphere.

### MVPA

We first classified the brain activities of the task block and those of the replay (observation only) block with leave one-session out cross-validation among the first 3 task sessions. A mixed-effects ANOVA was conducted on the classification accuracy in the precentral and postcentral cortex with the hemisphere (left/right) as a within-subject factor and the group (left-handed/right-handed) as a between-subject factor. The precentral cortex showed a significant interaction (*F*(1, 39) = 22.07, *p* < .001, *η_p_^2^* = .36) with no significant main effect of the group (*F*(1, 39) = 0.06, *p* = .81, *η_p_^2^* = .002) and the hemisphere (*F*(1, 39) = 1.01, *p* = .32, *η_p_^2^* = .025). Post-hoc analysis showed a significantly higher classification accuracy in the contralateral hemisphere for both the right-hand (*t*(20) = 4.46, *p* < .001, Cohen’s d = 1.00) and the left-hand group (*t*(19) = −2.36, *p* = .03, Cohen’s d = 0.55). The postcentral cortex also showed a significant interaction (*F*(1, 39) = 19.07, *p* < .001, *η_p_^2^* = .328) with no significant main effect of the group (*F*(1, 39) = 0.21, *p* = .65, *η_p_^2^* = .005) and the hemisphere (*F*(1, 39) = 0.007, *p* = .93, *η_p_^2^* = .0002). Post-hoc analysis showed a significantly higher classification accuracy in the contralateral hemisphere for both the right-hand group (*t*(20) = 3.41, *p* = .003, Cohen’s d = 0.73) and the left-hand group (*t*(19) = −2.81, *p* = .01, Cohen’s d = 0.64). These results thus showed successful classification of the task vs. non-task patterns, together with significantly higher accuracy in the contralateral sensorimotor cortex compared to the ipsilateral side of the hand used (Figure 4).

**Figure 4:**
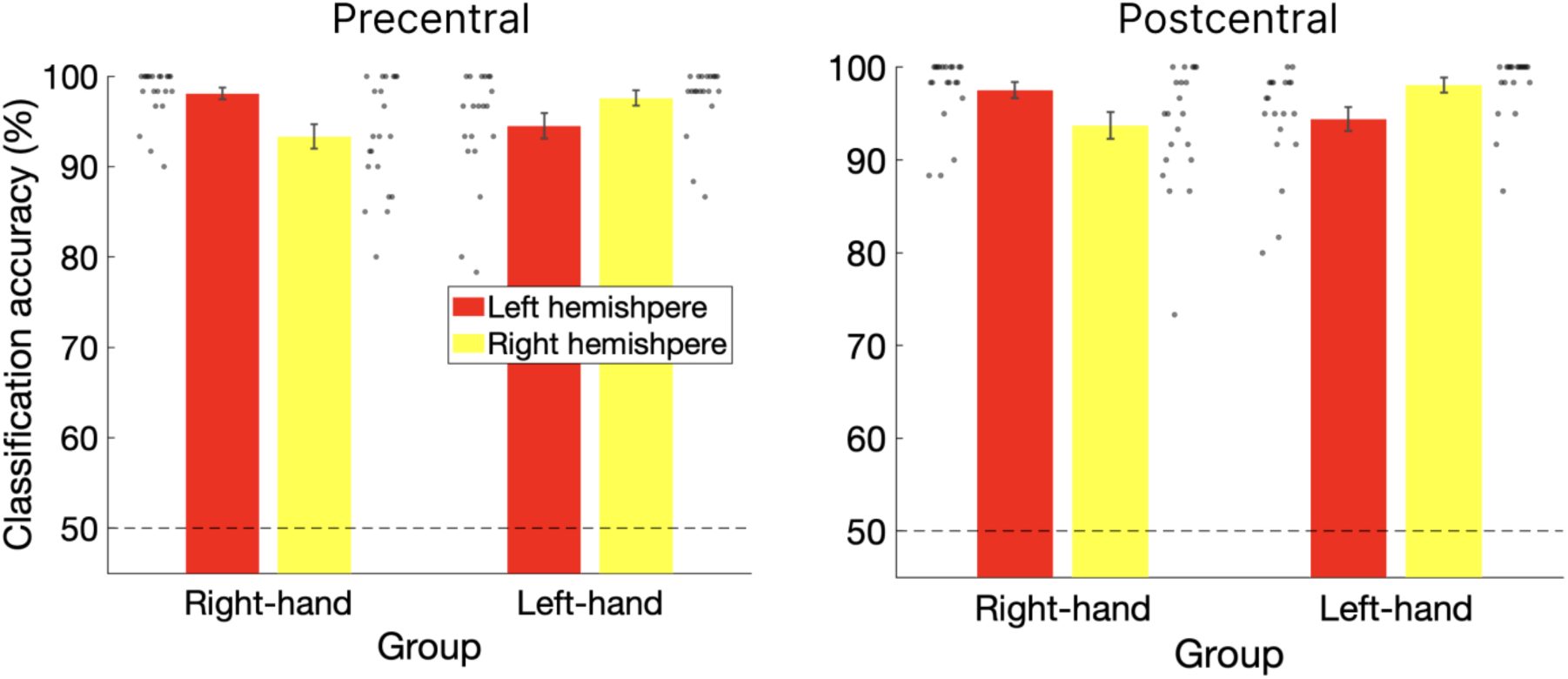
Results of the task and replay classification among task sessions The classification accuracy of task and replay (observation only) block activities among 3 task sessions in the precentral cortex (left) and postcentral cortex (right) for each group. The red and yellow bars showed the left and right hemispheres, respectively. The gray dots show the individual data. The black dotted lines denote the chance level of the classification. Error bars denote SEMs.

Next, we investigated whether the re-emergence of task-related activation patterns occurred more frequently after task training (post-RS) than before the task period (pre-RS). As in the previous analysis, the decoder was first trained to classify the brain activities between the task and the non-task patterns using all the 3 task sessions, and this decoder was tested with the activities of the pre- and post-RS to see whether a task-like brain activity pattern occurred during the resting-state session. A mixed-effects ANOVA was conducted on the frequency of occurrence of the volumes labeled as task patterns during pre-task and post-task resting-state sessions (pre-/post-RS) as a within-subject factor and the group (right-hand/left-hand) as a between-subject factor. The left precentral cortex showed a significant main effect of both the session (*F*(1, 39) = 4.52, *p* = .039, *η_p_^2^* = .10) and the group (*F*(1, 39) = 17.24, *p* < .001, *η_p_^2^* = .31) with a non-significant interaction (*F*(1, 39) = 1.78, *p* = .19, *η_p_^2^* = .04). This significant effect of the group without significant interaction suggests that the similar pattern of task activation was not only present in the post-RS but already in the pre-RS. The right precentral cortex showed a marginally significant main effect of the group (*F*(1, 39) = 4.02, *p* = .052, *η_p_^2^* = .094) with non-significant main effect of the session (*F*(1, 39) = 1.94, *p* = .17, *η_p_^2^* = .047) and a marginally significant interaction (*F*(1, 39) = 2.88, *p* = .10, *η_p_^2^* = .069). The left postcentral cortex showed a significant main effect of both the group (*F*(1, 39) = 5.61, *p* = .023, *η_p_^2^* = .13) and the session (*F*(1, 39) = 8.95, *p* = .005, *η_p_^2^* = .19) with a significant interaction (*F*(1, 39) = 4.64, *p* = .04, *η_p_^2^* = .11). The right postcentral cortex showed no significant main effect of both the group (*F*(1, 39) < .001, *p* = 1.00, *η_p_^2^* < .001) and the session (*F*(1, 39) = 2.63, *p* = .11, *η_p_^2^* = .06) with no significant interaction (*F*(1, 39) = 2.12, *p* = .15, *η_p_^2^* = .05) (Figure 5).

**Figure 5:**
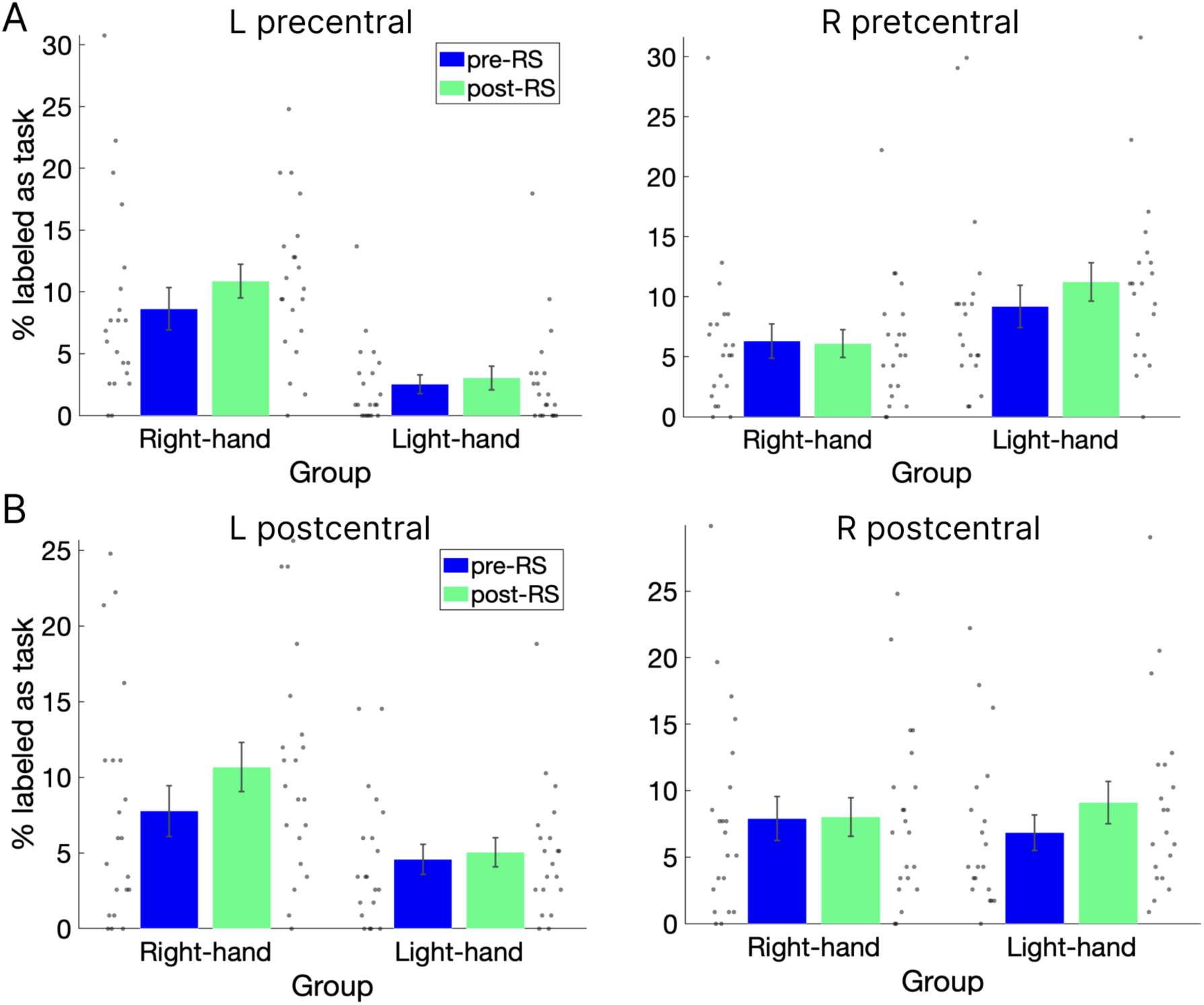
Percentage of the volumes labeled as task pattern during rest sessions. Percentage of the volumes in the precentral (A) and the postcentral cortex (B) classified as the task pattern compared with the non-task (observation only) pattern during the pre-RS (blue) and post-RS (green) for each group. The gray dots show the individual data. L, left hemisphere; R, right hemisphere. Error bars denote SEMs.

To directly compare differences in classification accuracy between pre- and post-task for each hemisphere and group, we subtracted the classification accuracy of pre-RS from post-RS using the same results. The precentral cortex showed a significant interaction (*F*(1, 39) = 5.93, *p* = .02, *η_p_^2^* = .12) with a non-significant main effect of both the groups (*F*(1, 39) = 0.07, *p* = .80, *η_p_^2^* < .001) and the hemisphere (*F*(1, 39) = .30, *p* = .59, *η_p_^2^* = .008). Post-hoc analysis showed larger numbers of task-labeled volumes in the contralateral hemisphere for both groups, and this difference is significant in the right-hand group (*t*(20) = 2.26, *p* = .04, Cohen’s d = 0.51) but not in the left-hand group (*t*(19) = −1.22, *p* = .24, Cohen’s d = 0.28). The postcentral cortex showed a significant interaction (*F*(1, 39) = 8.79, *p* = .001, *η_p_^2^* = .18) with no significant main effect of the group (*F*(1, 39) = 0,02, *p* = .90 *η_p_^2^* < .001) and the hemisphere (*F*(1, 39) = 0,40, *p* = .53, *η_p_^2^* = .01). Post-hoc analysis showed significantly or marginally significantly larger numbers of task-labeled volumes in the contralateral hemisphere for both the right-hand group ((*t*(20) = 2.28, *p* = .03, Cohen’s d = 0.51) and the left-hand group ((*t*(19) = −1.94, *p* = .07, Cohen’s d = 0.44). Our results thus showed a higher frequency of reactivations in the primary sensorimotor cortex during the post-learning compared to the pre-learning period, and that this effect is specific to the primary sensorimotor cortex contralateral to the hand used (Figure 6A).

**Figure 6:**
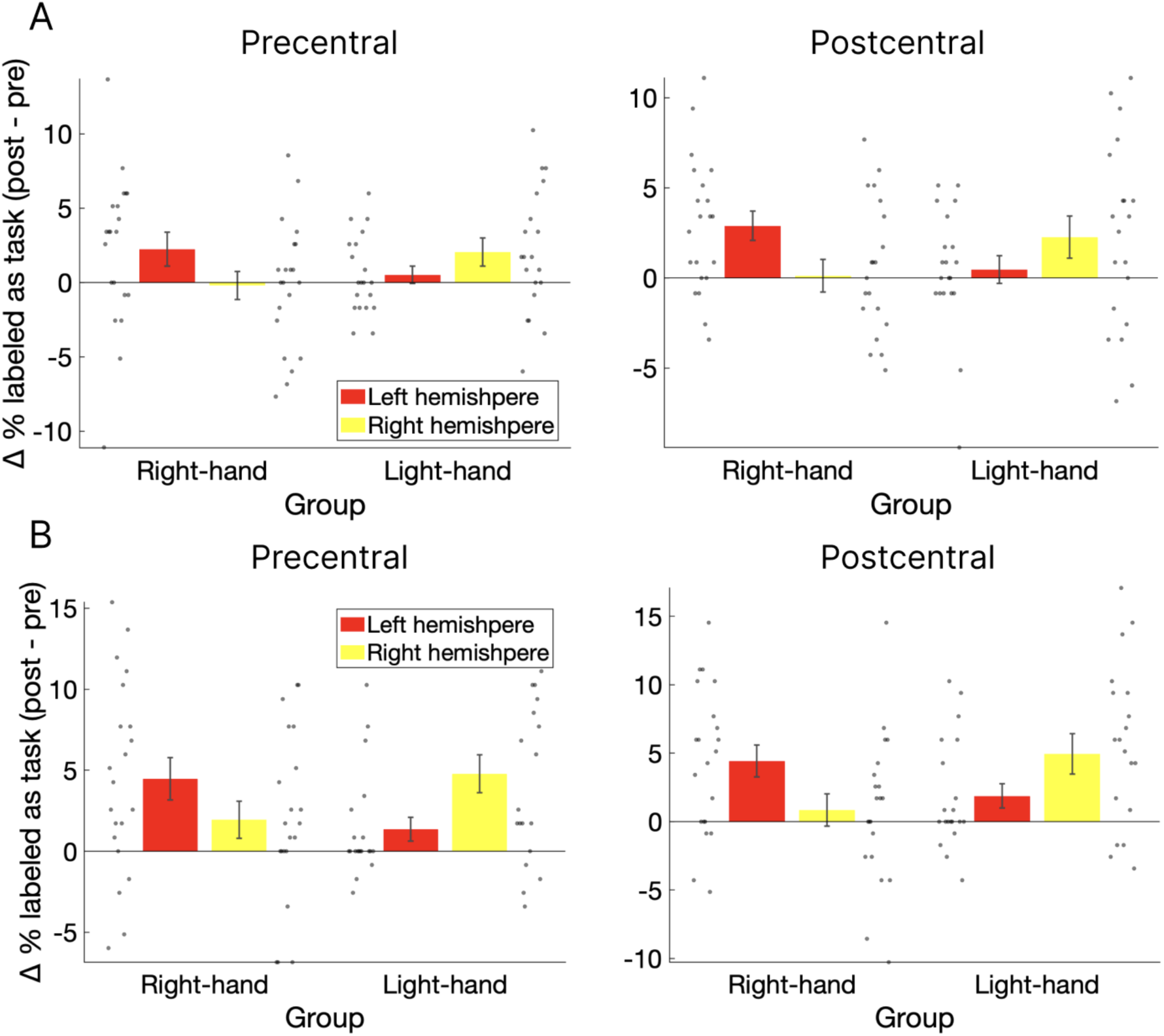
Difference in the frequency of the volumes labeled as the task in pre-RS and post-RS The difference in frequency between pre-RS and post-RS volumes labeled as (A) or more similar to (B) the task pattern compared with the non-task (observation only) pattern in the precentral and the postcentral cortex with the classification analysis (A) and the RSA (B). The red and yellow bars show the left and right hemispheres, respectively. The gray dots show the individual data. Error bars denote SEMs.

We further conducted the same analysis with the RSA results. The precentral cortex showed a significant interaction (*F*(1, 39) = 10.23, *p* = .003, *η_p_^2^* = .21) with no significant main effect of both the group (*F*(1, 39) = 0.01, *p* = .91, *η_p_^2^* < .001) and the hemisphere (*F*(1, 39) = .23, *p* = .63, *η_p_^2^* = .005). Post-hoc analysis showed significantly or marginally significantly larger numbers of task-labeled volumes in the contralateral hemisphere for both the right-hand group ((*t*(20) = 1.75, *p* = .10, Cohen’s d = 0.39) and the left-hand group ((*t*(19) = −3.00, *p* = .008, Cohen’s d = 068). The postcentral cortex also showed a significant interaction (*F*(1, 39) = 11.86, *p* = .001, *η_p_^2^* = .23) with no significant main effect of the group (*F*(1, 39) = 0.31, *p* = .58, *η_p_^2^* = .008) and the hemisphere (*F*(1, 39) = 0.07, *p* = .80, *η_p_^2^* = .002). Post-hoc analysis showed significantly larger numbers of task-labeled volumes in the contralateral hemisphere for both the right-hand group ((*t*(20) = 2.37, *p* = .03, Cohen’s d = 0.53) and the left-hand group ((*t*(19) = −2.59, *p* = .02, Cohen’s d = 059). These results are consistent with the previous classification analysis, regarding the higher frequency of reactivations during the post-learning in the primary sensorimotor cortex contralateral to the hand used (Figure 6B).

Finally, we investigated the relationship between the frequency of reactivation during the rest and behavioral performance. We thus analyzed the correlation between the increase in the percentage of labeled task patterns from pre-RS to post-RS in the contralateral hemisphere of each group and the decrease in tracking errors from the average of the first 3 task sessions to the last task session. The results of the right-hand group showed a significant correlation in the left postcentral cortex (*R* = .64, *p_Bonf_* < .05: Figure 7C), but no significant correlation in the left precentral cortex (*R* = −.04, *p_Bonf_* > .05: Figure 7A). For the left-hand group, no significant correlation was found in either the right precentral (*R* = −.14, *p_Bonf_* > .05: Figure 7B) or the postcentral cortex (*R* = −.48, *p_Bonf_* > .05: Figure 7D).

**Figure 7:**
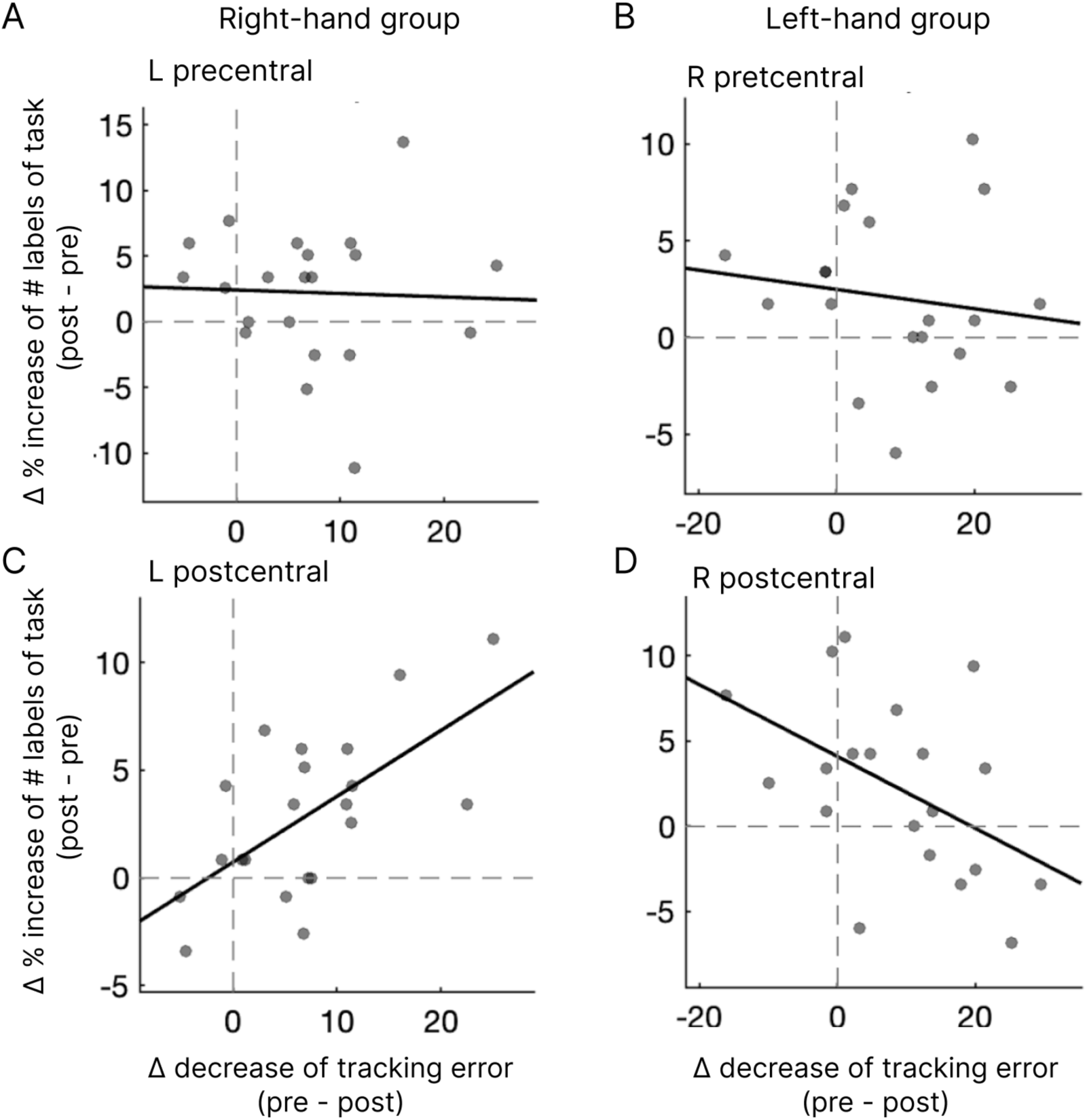
Correlation between MVPA results and behavioral performance The correlation between the increased percentage of labeled task patterns and the decreased tracking error. The increase indicated the percentage difference between pre-RS and post-RS labeling as task patterns in the contralateral precentral and the postcentral cortex. L, left hemisphere; R, right hemisphere.

## Discussion

In this study, we used fMRI with MVPA to reveal the re-emergence of task-related activation patterns in the resting state after the visuomotor learning task. We first classified the brain activities of the task and those of the replay (observation only) using the first 3 task sessions, which showed successful classification with significantly higher accuracy in the contralateral sensorimotor cortex compared to the ipsilateral side of the hand used. Next, we investigated whether the reappearance of task-related activation patterns occurred more frequently after task training (post-RS) than before the task period (pre-RS). Both multivariate classification and RSA consistently revealed a significantly higher number of reactivations in the primary sensorimotor cortex during the post-learning period compared to the pre-learning period. Furthermore, this effect is specific to the primary sensorimotor cortex contralateral to the hand used and is significantly correlated with motor performance after rest. Our findings reveal the reactivation of task-related patterns in the primary sensorimotor cortex during offline visuomotor learning.

A number of previous studies using spatial navigation tasks in rodents have shown that patterns of neural activity associated with spatial experience are replayed in the hippocampus during rest, which is related to memory consolidation (Euston et al., 2007; Carr et al., 2011; Grosmark and Buzsáki, 2016). In addition to spatial navigation tasks, motor learning also induces replay-like activations after rodents learn a skilled upper-limb task, which is related to offline improvements in motor performance (Gulati et al., 2014; Ramanathan et al., 2015). A recent human study with magnetoencephalography (MEG) also revealed fast waking neural replay during the same rest periods in which rapid consolidation occurs across the hippocampus and neocortex after learning of novel motor sequence (Buch et al., 2021). A human fMRI study reported that the multi-voxel pattern in resting-state brain activity corresponds to either wrist or finger movements in the motor-related areas of each hemisphere of the cerebrum and cerebellum (Kusano et al., 2021), which could possibly acquired through prior experience of bodily movements. Another human fMRI study also reported a replay activity in the hippocampus during wakeful rest after decision-making with non-spatial sequences (Schuck and Niv, 2019), which indicates that reactivation is a more domain-general process, not limited to specific cognitive functions.

A recent invasive brain recording study measured the activity of intracortical microelectrode arrays in the left anterior tegmental gyrus while the participants were completing a novel motor task and a subsequent night’s sleep to determine whether replay occurs after motor learning (Rubin et al., 2022). They found that neural signals recorded overnight replayed the target sequences in the memory game at a significantly higher frequency than chance. Another human intracranial recording study also revealed a replay of activity for a learned motor sequence during awake rest in the motor cortex with a comparable rate of replay event (Eichenlaub et al., 2020). These invasive brain recording studies indicate an offline replay of neural firing patterns that underlie the waking experience and play a role in memory consolidation.

A previous study using non-invasive brain stimulation showed that when the primary motor cortex is disrupted by the transcranial magnetic stimulation (TMS) following motor learning, there is a reduction in learning, and this effect is specific to the waking rest period and is not observed during sleep (Robertson et al., 2005). Another TMS study also showed disrupting the primary motor cortex impairs early boost but not delayed gains in performance in motor sequence learning (Hotermans et al., 2008). These previous brain stimulation studies indicate the roles of the primary motor cortex for offline learning of visuomotor control.

Regarding the relationship between reactivation and behavior, we found a significant correlation between the frequency of reactivation patterns and the decrease in tracking errors from pre- to post-learning in the left postcentral cortex (Figure 7C), but not in the precentral cortex (Figure 7A), for the right-hand group. This finding that reactivation in the primary somatosensory cortex correlates with behavioral performance may suggest that reactivation at the sensory level, rather than the motor level, is more important for later performance improvement. In contrast, we found no significant correlation in both the precentral and postcentral cortex for the left-hand group (Figure 7B & D). As all of the participants are right-handed, this effect might be dependent on the handedness. The left-hand group also showed a non-significant or marginally significant increase in the task-like volumes during post-RS compared with pre-RS in the precentral and postcentral cortex, respectively (Figure 6A). As the left-hand group showed significantly more tracking errors compared with the right-hand group (Figure 2), insufficient learning with the non-dominant hand may have affected the less clear effects of the left-hand group, which need further investigation.

The present experiment showed that the contralateral sensorimotor cortex resembles task activity compared to the ipsilateral side, not only in the post-learning but also in the pre-learning resting period. The fact that this task-like activity was already present before the task is somewhat puzzling, but this may be related to the “pre-play” reported in the spontaneous activity of rodents. Pre-play, as opposed to replay, is a phenomenon in which hippocampal neurons are activated sequentially according to the place fields when rodents perform a spatial navigation task, but this occurs during rest before the animal actually performs the task (Dragoi and Tonegawa, 2011, 2013). This phenomenon suggests that hippocampal activation during rest may function not only in memory consolidation and retrieval but also in the planning stage. A recent human fMRI study also reported preplay-like activations for the acquisition of new semantic knowledge (Kurashige et al., 2018). Considering these previous studies, the current finding may also reflect preplay-like activations for learning visuomotor skills, which should be investigated in the future.

The current study has some limitations. Firstly, the increased activity pattern similar to the task during the post-learning resting period may simply reflect residual brain activity after performing the motor task. We consider this possibility unlikely for the following reasons. We displayed the frequency of labels that were classified as the task for the classification analysis and the RSA during the resting periods (Supplementary Figure 1). If the observed effect reflects simply residual brain activity after a motor task, we should see more task-like activity patterns immediately after the task is completed. Instead, the results showed that the task-assigned labels were distributed throughout the 6-minute resting period. This suggests that this is not residual task-related activity, but rather sustained brain activity during the resting state. Secondly, related to the previous point, the observed reactivation may reflect that the subjects were intentionally or unintentionally rehearsing the previous motor learning experience at rest even though the subjects were asked to remain rested (see Methods). We consider this possibility unlikely, because also in this case, the residual effects of the motor memory should gradually diminish over time. As was shown above, the task-like volume did not decrease over time, so we thus consider it unlikely that the observed reactivation was a rehearsal of motor memory. However, further experiments, such as imposing different kinds of cognitively demanding tasks to prevent such rehearsal at rest, will be needed in the future to clearly reject this possibility. Thirdly, the present study deals only with one rotational transformation and not with brain activity specific to a particular transformation rule. Our previous study has shown that multiple rotational transformations are acquired in the primary sensorimotor cortex (Ogawa and Imamizu, 2013), and the relevance of specific motor skills to the reproduction of brain activity patterns is unclear in the present experiment. Lastly, the present experiment only deals with kinematic motor adaptation with a rotational transformation, and it is not clear whether the results can be generalized to other motor adaptations and learning tasks, including sequential motor learning (e.g., serial reaction time task; SRTT) (Robertson, 2007). This should also be investigated in future studies.

In summary, we found a significant increase in task-related activities in the post-learning period compared with the pre-learning period. In addition, this effect is specific to the primary sensorimotor cortex contralateral to the hand used and significantly correlated with motor improvement after rest. Our finding revealed the reactivation of task-related patterns in the primary sensorimotor cortex for visuomotor learning.

## Supplementary Figure

**Supplementary Figure 1:**
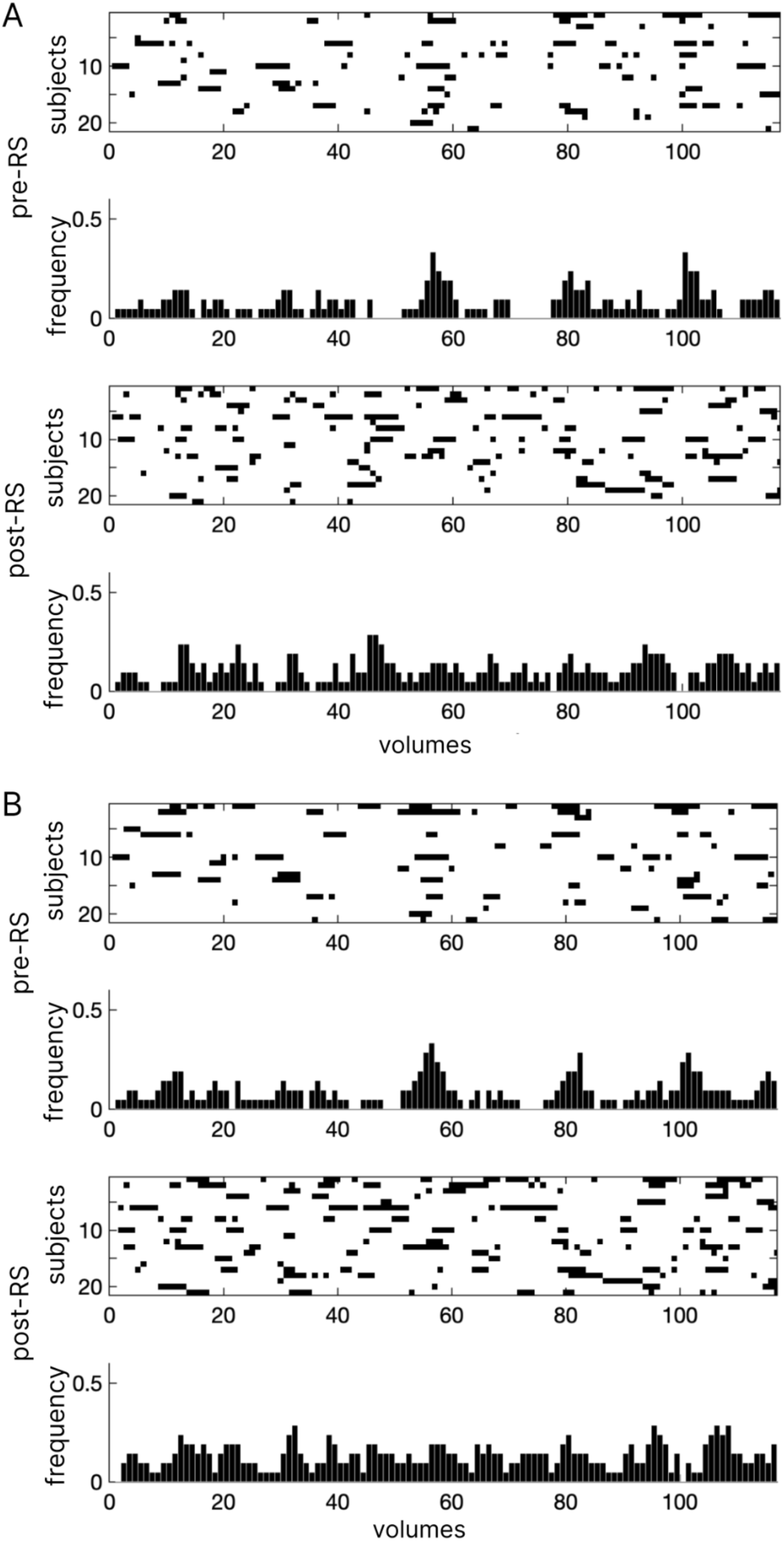
Example of MVPA decoded labels in the left postcentral cortex of the right-hand group The frequency of spatiotemporal patterns labeled as task patterns during rest sessions using the MVPA classification analysis (A) and the RSA (B). The black label indicates the volume classified as (A) or more similar (B) to the task pattern compared with the non-task (observation only) pattern. The upper two figures in each A and B show the distribution (upper) and frequency (lower) during the pre-RS, and the lower two figures show those during the post-RS.

## Acknowledgments

This work was supported by JSPS KAKENHI Grant Numbers JP17H04683 & JP21H00958 to K.O.

## Notes

The authors declare no conflict of interest associated with this manuscript.

### Competing Interest Statement

The authors have declared no competing interest.

### Summary of Updates

The labels in Figure 3 were incorrect and have been corrected.

## References

Albert NB, Robertson EM, Miall RC (2009) The resting human brain and motor learning. Curr Biol 19:1023–1027.

Berkes P, Orbán G, Lengyel M, Fiser J (2011) Spontaneous cortical activity reveals hallmarks of an optimal internal model of the environment. Science 331:83–87.

Brashers-Krug T, Shadmehr R, Bizzi E (1996) Consolidation in human motor memory. Nature 382:252–255.

Buch ER, Claudino L, Quentin R, Bönstrup M, Cohen LG (2021) Consolidation of human skill linked to waking hippocampo-neocortical replay. Cell Rep 35:109193.

Carr MF, Jadhav SP, Frank LM (2011) Hippocampal replay in the awake state: a potential substrate for memory consolidation and retrieval. Nat Neurosci 14:147–153.

Cepeda NJ, Pashler H, Vul E, Wixted JT, Rohrer D (2006) Distributed practice in verbal recall tasks: A review and quantitative synthesis. Psychol Bull 132:354–380.

Cohen DA, Pascual-Leone A, Press DZ, Robertson EM (2005) Off-line learning of motor skill memory: a double dissociation of goal and movement. Proc Natl Acad Sci U S A 102:18237–18241.

Dragoi G, Tonegawa S (2011) Preplay of future place cell sequences by hippocampal cellular assemblies. Nature 469:397–401.

Dragoi G, Tonegawa S (2013) Distinct preplay of multiple novel spatial experiences in the rat. Proc Natl Acad Sci U S A 110:9100–9105.

Eichenlaub J-B, Jarosiewicz B, Saab J, Franco B, Kelemen J, Halgren E, Hochberg LR, Cash SS (2020) Replay of Learned Neural Firing Sequences during Rest in Human Motor Cortex. Cell Rep 31:107581.

Euston DR, Tatsuno M, McNaughton BL (2007) Fast-forward playback of recent memory sequences in prefrontal cortex during sleep. Science 318:1147–1150.

Fox MD, Raichle ME (2007) Spontaneous fluctuations in brain activity observed with functional magnetic resonance imaging. Nat Rev Neurosci 8:700–711.

Gentili RJ, Bradberry TJ, Oh H, Costanzo ME, Kerick SE, Contreras-Vidal JL, Hatfield BD (2015) Evolution of cerebral cortico-cortical communication during visuomotor adaptation to a cognitive-motor executive challenge. Biol Psychol 105:51–65.

Gerbier E, Toppino TC, Koenig O (2015) Optimising retention through multiple study opportunities over days: The benefit of an expanding schedule of repetitions. Memory 23:943–954.

Gregory MD, Agam Y, Selvadurai C, Nagy A, Vangel M, Tucker M, Robertson EM, Stickgold R, Manoach DS (2014) Resting state connectivity immediately following learning correlates with subsequent sleep-dependent enhancement of motor task performance. Neuroimage 102:666–673.

Grosmark AD, Buzsáki G (2016) Diversity in neural firing dynamics supports both rigid and learned hippocampal sequences. Science 351:1440–1443.

Guidotti R, Del Gratta C, Baldassarre A, Romani GL, Corbetta M (2015) Visual Learning Induces Changes in Resting-State fMRI Multivariate Pattern of Information. J Neurosci 35:9786–9798.

Gulati T, Ramanathan DS, Wong CC, Ganguly K (2014) Reactivation of emergent task-related ensembles during slow-wave sleep after neuroprosthetic learning. Nat Neurosci 17:1107– 1113.

Han F, Caporale N, Dan Y (2008) Reverberation of recent visual experience in spontaneous cortical waves. Neuron 60:321–327.

Hatta T, Nakatsuka Z (1975) Handedness inventory. In: Papers on Celebrating 63rd Birthday of Prof. Ohnishi, pp 224–245. Osaka City University.

Hotermans C, Peigneux P, de Noordhout AM, Moonen G, Maquet P (2008) Repetitive transcranial magnetic stimulation over the primary motor cortex disrupts early boost but not delayed gains in performance in motor sequence learning. Eur J Neurosci 28:1216– 1221.

Imamizu H, Miyauchi S, Tamada T, Sasaki Y, Takino R, Pütz B, Yoshioka T, Kawato M (2000) Human cerebellar activity reflecting an acquired internal model of a new tool. Nature 403:192–195.

Ji D, Wilson MA (2007) Coordinated memory replay in the visual cortex and hippocampus during sleep. Nat Neurosci 10:100–107.

Kamitani Y, Sawahata Y (2010) Spatial smoothing hurts localization but not information: Pitfalls for brain mappers. NeuroImage 49:1949–1952 Available at: 10.1016/j.neuroimage.2009.06.040.

Klinzing JG, Niethard N, Born J (2019) Mechanisms of systems memory consolidation during sleep. Nat Neurosci 22:1598–1610.

Kornmeier J, Sosic-Vasic Z (2012) Parallels between spacing effects during behavioral and cellular learning. Front Hum Neurosci 6:203.

Kriegeskorte N, Mur M, Bandettini P (2008) Representational similarity analysis - connecting the branches of systems neuroscience. Front Syst Neurosci 2:1–28.

Kurashige H, Yamashita Y, Hanakawa T, Honda M (2018) A Knowledge-Based Arrangement of Prototypical Neural Representation Prior to Experience Contributes to Selectivity in Upcoming Knowledge Acquisition. Front Hum Neurosci 12:111.

Kurth-Nelson Z, Behrens T, Wayne G, Miller K, Luettgau L, Dolan R, Liu Y, Schwartenbeck P (2023) Replay and compositional computation. Neuron 111:454–469.

Kusano T, Kurashige H, Nambu I, Moriguchi Y, Hanakawa T, Wada Y, Osu R (2021) Wrist and finger motor representations embedded in the cerebral and cerebellar resting-state activation. Brain Struct Funct Available at: 10.1007/s00429-021-02330-8.

Lewis CM, Baldassarre A, Committeri G, Romani GL, Corbetta M (2009) Learning sculpts the spontaneous activity of the resting human brain. Proceedings of the National Academy of Sciences 106:17558–17563.

Lin CH., Yang HC, Ye YL, Huang SL, Chiang MC, Knowlton BJ, Wu AD, Iacoboni M (2018) Contextual interference enhances motor learning through increased resting brain connectivity during memory consolidation. NeuroImage.

Manuel AL, Guggisberg AG, Thézé R, Turri F, Schnider A (2018) Resting-state connectivity predicts visuo-motor skill learning. Neuroimage 176:446–453.

Mur M, Bandettini PA, Kriegeskorte N (2009) Revealing representational content with pattern-information fMRI - An introductory guide. Soc Cogn Affect Neurosci 4:101–109.

Ogawa K, Imamizu H (2013) Human sensorimotor cortex represents conflicting visuomotor mappings. J Neurosci 33:6412–6422.

Ogawa K, Mitsui K, Imai F, Nishida S (2019) Long-term training-dependent representation of individual finger movements in the primary motor cortex. Neuroimage 202:116051.

Oldfield RC (1971) The assessment and analysis of handedness: the Edinburgh inventory. Neuropsychologia 9:97–113.

Pavlides C, Winson J (1989) Influences of hippocampal place cell firing in the awake state on the activity of these cells during subsequent sleep episodes. J Neurosci 9:2907–2918.

Press DZ, Casement MD, Pascual-Leone A, Robertson EM (2005) The time course of off-line motor sequence learning. Cognitive Brain Research 25:375–378.

Raichle ME, MacLeod AM, Snyder AZ, Powers WJ, Gusnard DA, Shulman GL (2001) A default mode of brain function. Proc Natl Acad Sci U S A 98:676–682.

Ramanathan DS, Gulati T, Ganguly K (2015) Sleep-Dependent Reactivation of Ensembles in Motor Cortex Promotes Skill Consolidation. PLoS Biol 13:e1002263.

Robertson EM (2007) The Serial Reaction Time Task: Implicit Motor Skill Learning? Journal of Neuroscience 27:10073–10075.

Robertson EM, Pascual-Leone A, Miall RC (2004) Current concepts in procedural consolidation. Nat Rev Neurosci 5:576–582.

Robertson EM, Press DZ, Pascual-Leone A (2005) Off-line learning and the primary motor cortex. J Neurosci 25:6372–6378.

Rubin DB, Hosman T, Kelemen JN, Kapitonava A, Willett FR, Coughlin BF, Halgren E, Kimchi EY, Williams ZM, Simeral JD, Hochberg LR, Cash SS (2022) Learned Motor Patterns Are Replayed in Human Motor Cortex during Sleep. J Neurosci 42:5007–5020.

Sami S, Robertson EM, Miall RC (2014) The Time Course of Task-Specific Memory Consolidation Effects in Resting State Networks. Journal of Neuroscience 34:3982–3992.

Schuck NW, Niv Y (2019) Sequential replay of nonspatial task states in the human hippocampus. Science 364 Available at: 10.1126/science.aaw5181.

Shadmehr R, Brashers-Krug T (1997) Functional stages in the formation of human long-term motor memory. J Neurosci 17:409–419.

Tambini A, Davachi L (2019) Awake Reactivation of Prior Experiences Consolidates Memories and Biases Cognition. Trends Cogn Sci 23:876–890.

Tambini A, Ketz N, Davachi L (2010) Enhanced Brain Correlations during Rest Are Related to Memory for Recent Experiences. Neuron 65:280–290.

Tzourio-Mazoyer N, Landeau B, Papathanassiou D, Crivello F, Etard O, Delcroix N, Mazoyer B, Joliot M (2002) Automated anatomical labeling of activations in SPM using a macroscopic anatomical parcellation of the MNI MRI single-subject brain. Neuroimage 15:273–289.

Vahdat S, Darainy M, Milner TE, Ostry DJ (2011) Functionally specific changes in resting-state sensorimotor networks after motor learning. J Neurosci 31:16907–16915.

Wolpert DM, Ghahramani Z (2000) Computational principles of movement neuroscience. Nature 3 Available at: 10.1038/81497.

Wu J, Srinivasan R, Kaur A, Cramer SC (2014) Resting-state cortical connectivity predicts motor skill acquisition. Neuroimage 91:84–90.

